# Alterations in grey matter structure linked to frequency-specific cortico-subcortical connectivity in schizophrenia via multimodal data fusion

**DOI:** 10.1101/2023.07.05.547840

**Authors:** Marlena Duda, Ashkan Faghiri, Aysenil Belger, Juan R. Bustillo, Judith M. Ford, Daniel H. Mathalon, Bryon A. Mueller, Godfrey D. Pearlson, Steven G. Potkin, Adrian Preda, Jing Sui, Theo G.M. Van Erp, Vince D. Calhoun

## Abstract

Schizophrenia (SZ) is a complex psychiatric disorder that is currently defined by symptomatic and behavioral, rather than biological, criteria. Neuroimaging is an appealing avenue for SZ biomarker development, as several neuroimaging-based studies comparing individuals with SZ to healthy controls (HC) have shown measurable group differences in brain structure, as well as functional brain alterations in both static and dynamic functional network connectivity (sFNC and dFNC, respectively). The recently proposed filter-banked connectivity (FBC) method extends the standard dFNC sliding-window approach to estimate FNC within an arbitrary number of distinct frequency bands. The initial implementation used a set of filters spanning the full connectivity spectral range, providing a unified approach to examine both sFNC and dFNC in a single analysis. Initial FBC results found that individuals with SZ spend more time in a less structured, more disconnected low-frequency (i.e., static) FNC state than HC, as well as preferential SZ occupancy in high-frequency connectivity states, suggesting a frequency-specific component underpinning the functional dysconnectivity observed in SZ. Building on these findings, we sought to link such frequency-specific patterns of FNC to covarying data-driven structural brain networks in the context of SZ. Specifically, we employ a multi-set canonical correlation analysis + joint independent components analysis (mCCA + jICA) data fusion framework to study the connection between grey matter volume (GMV) maps and FBC states across the full connectivity frequency spectrum. Our multimodal analysis identified two joint sources that captured co-varying patterns of frequency-specific functional connectivity and alterations in GMV with significant group differences in loading parameters between the SZ group and HC. The first joint source linked frequency-modulated connections between the subcortical and sensorimotor networks and GMV alterations in the frontal and temporal lobes, while the second joint source identified a relationship between low-frequency cerebellar-sensorimotor connectivity and structural changes in both the cerebellum and motor cortex. Together, these results show a strong connection between cortico-subcortical functional connectivity at both high and low frequencies and alterations in cortical GMV that may be relevant to the pathogenesis and pathophysiology of SZ.

## 1 Introduction

Neuroimaging has become a valuable tool for noninvasively studying the human brain. Several neuroimaging tools now exist that are capable of capturing brain structure and tissue type at various anatomical levels (e.g., structural MRI [sMRI] and diffusion MRI [dMRI]), as well as indirectly estimating brain function or activity through characteristic source signals of the underlying neuronal, metabolic, or hemodynamic activity (e.g., electroencephalography/ magnetoencephalography [EEG/MEG], positron emission tomography [PET], functional MRI [fMRI], respectively). While each of these imaging modalities is powerful and useful in its own right, each provides a unique yet incomplete picture of the brain. Furthermore, each modality is accompanied by its own inherent limitations on spatial and temporal resolution, imposed by the technical specifications of each image acquisition type. To gain a more complete picture of an individual’s neural landscape and overcome the limitations of any single imaging modality, multimodal analyses can be utilized to combine and leverage the rich and complementary information available across various neuroimaging types.

Multimodal data fusion represents a class of analytical approaches that aim to integrate data across complementary neuroimaging modalities. Simpler approaches to data fusion may connect results from separate unimodal analyses through post-hoc correlations or use the results from one modality to constrain the model for another modality (i.e., asymmetric data fusion). Such multimodal approaches can provide useful insights but ultimately do not take full advantage of the available joint (i.e., cross-modal) information, which is the key aim of the so-called "symmetric” multimodal fusion approaches (Calhoun & Sui, 2016; Sui et al., 2012). This family of data fusion approaches considers each imaging modality equally to estimate a final joint result and can be further broken down into two categories: model-based vs. data-driven approaches. While model-based approaches can be valuable when there is sufficient a priori knowledge about the problem being studied, data-driven fusion approaches are often advantageous because they impose fewer assumptions on the interrelationships between the data types and enable exploration of the entire voxel space rather than limiting to only those interrelationships that were explicitly modeled prior. For this reason, data-driven approaches are especially useful for studying complex psychiatric disorders such as schizophrenia, where there is still much to be learned about the etiology (Ayano, 2016; Misiak et al., 2018).

Existing data-driven approaches often use blind or semi-blind variations of linear mixture models to reveal hidden linkages between feature spaces derived from two or more imaging modalities. These approaches include, but are not limited to, joint independent component analysis (jICA) (Calhoun et al., 2006), linked ICA (Groves et al., 2011), partial least squares (PLS) (Martínez-Montes et al., 2004), and multimodal/multiset canonical correlation analysis (mCCA) (Correa et al., 2007, 2010) for blind approaches, and coefficient constrained ICA (cc-ICA) (Sui et al., 2009) and parallel ICA (Liu et al., 2009) for semi-blind approaches. Each of these multivariate approaches differ in their optimization procedures and basic limitations, but just as multimodal analyses can combine complementary data types to overcome the limitations of each, combining multiple multivariate fusion algorithms has been shown to mitigate the limiting effects of the individual methods (Sui et al., 2011).

One example of a combined approach is the mCCA + jICA fusion framework (Sui et al., 2011, 2013). In jICA the objective is to estimate sources that are maximally independent from one another, but the shared mixing matrix across the datasets assumes a strong correlation between the distinct modalities. Conversely, mCCA maximizes the correlations of inter-subject mixing profiles, thus allowing for varying correlations between the joint sources, but may result in spatial maps for the joint sources that are not sufficiently different from one another. However, the combined mCCA + jICA model is designed to allow for the identification of both strongly and weakly correlated joint components that are also independent from one another by employing mCCA in the first step to generate flexible linkages between the modalities and subsequently applying jICA on the associated maps in the second step.

The mCCA + jICA framework has been utilized for several neuroimaging data fusion studies of complex disorders, including schizophrenia (SZ). SZ is a chronic and debilitating neuropsychiatric syndrome marked by a variety of mental and behavioral symptoms including positive symptoms such as delusions, hallucinations, disorganized speech and/or behavior, negative symptoms such as diminished emotional expression and avolition, and cognitive deficits impacting on an individual’s professional life and interpersonal relationships (American Psychiatric Association, 2013). There is considerable evidence that functional, structural, genetic, and epigenetic alterations are associated with SZ; however, none yet have proven to be sufficiently reliable for use as clinical biomarkers, especially at an individual level (Fornito et al., 2012; Khavari & Cairns, 2020; Kraguljac et al., 2021; Pantelis et al., 2009; Pickard, 2015; Rodrigues-Amorim et al., 2017). While this can be due to the substantial heterogeneity of SZ and imperfections in current defining diagnostic criteria, it has also been suggested that this lack of clinically relevant diagnostic markers can be attributed, at least in part, to the oversaturation of unimodal analyses and the lack of effective multimodal studies, thus missing important neurobiological components of SZ that can only be partially detected by individual modalities (Calhoun & Sui, 2016). As the importance of multimodal fusion analyses continues to be recognized, the number of multimodal studies of SZ has increased, the results of which show evidence for strong linkages between structural, functional, and even genetic factors of the disease (Acar et al., 2019; DeRamus et al., 2022; Lottman et al., 2018; Y. Zhang et al., 2022).

The increasing interest in studying “time evolving” or dynamic FNC and how these dynamics may relate to psychiatric syndromes like SZ has begun to be incorporated into multimodal studies of disease (Abrol et al., 2017; Calhoun et al., 2014). Currently, dFNC is the object of much debate in the field. However, much of the skepticism surrounding dFNC is based on the embedded assumptions of the common sliding window Pearson correlation (SWPC), namely issues with assuming the timescale of the dynamics by imposing a static and somewhat arbitrarily chosen window size (Hindriks et al., 2016; Shakil et al., 2018), resulting in a low-pass filtered view of the connectivity time series (Hutchison et al., 2013; Leonardi & Van De Ville, 2015; Sakoğlu et al., 2010; Thompson & Fransson, 2015). A recent method termed filter-banked connectivity (FBC) extends the SWPC and provides a unified approach for estimating FNC that includes the information of both static and dynamic FNC simultaneously (Faghiri et al., 2021). Furthermore, by employing frequency-tiling (i.e., decomposition of the original signal within various frequency ranges) via filter banks the FBC enables estimation of changing FNC in specified frequency bands, effectively providing estimates of dFNC at various timescales in a single approach. What distinguishes the FBC from other frequency-based dFNC approaches that have been implemented in the past (e.g., cross wavelet coherence (Chang & Glover, 2010; Yaesoubi et al., 2015)) is that the frequency tiling occurs directly in the connectivity domain, rather than in the functional activity domain. This detail is key because the relationship between the activation and connectivity domains is possibly non-linear, and since the final inference is based on connectivity it is critical that all frequency tiling steps be also performed in the connectivity domain to prevent misinterpretation of the frequency information. Initial results demonstrated that FBC was indeed capable of identifying dFNC states in high-frequency ranges that were missed by SWPC (Faghiri et al., 2021). Further analysis of a SZ and control cohort with the FBC approach identified a relatively unstructured and disconnected low-frequency (i.e., close to static) FNC state predominantly occupied by SZ subjects, in contrast to an organized and highly connected low-frequency state that was predominantly occupied by controls. This study also showed preferential SZ occupancy in high-frequency connectivity states (Faghiri et al., 2021). These results are consistent with previous frequency-based studies of the activity domain that reported higher power at higher frequencies in individuals with SZ compared to controls (Alonso-Solís et al., 2017; Calhoun et al., 2011; Garrity et al., 2007); however care must be taken when comparing results from the activity vs. connectivity domain analyses. Taken together, these results suggest there may exist an important frequency-specific functional component underpinning the pathophysiology of SZ.

Here, we sought to extend this line of work by investigating the relationship between frequency-specific functional connectivity patterns and structural brain features that are associated with SZ. Specifically, we link frequency-specific connectivity states derived with FBC to sMRI grey matter volume (GMV) maps using the mCCA + jICA framework introduced above. Through this work we aim to further uncover the role that both slow (low-frequency) and rapid (high-frequency) changes in FNC may play in the pathophysiology of SZ by identifying group-discriminative structure-function relationships that exist within distinct spectral ranges.

## 2 Methods

### 2.1 Data Description

We utilized an age- and gender-matched dataset (Keator et al., 2016) including 310 individuals, 150 with SZ (114 male, avg. age = 38.8 years) and 160 healthy controls (HC; 115 male, avg. age = 37.0 years) that met our subject inclusion criteria of high-quality registration to EPI template and head motion translation of less than 3° rotation and 3 mm translation in all directions (Fu et al., 2021). Informed consent was obtained from each participant prior to MRI scanning and all studies were approved by the Institutional Review Boards of institutions involved in data collection (Keator et al., 2016). Detailed demographics of the SZ group are presented in Table 1.

**Table 1.**
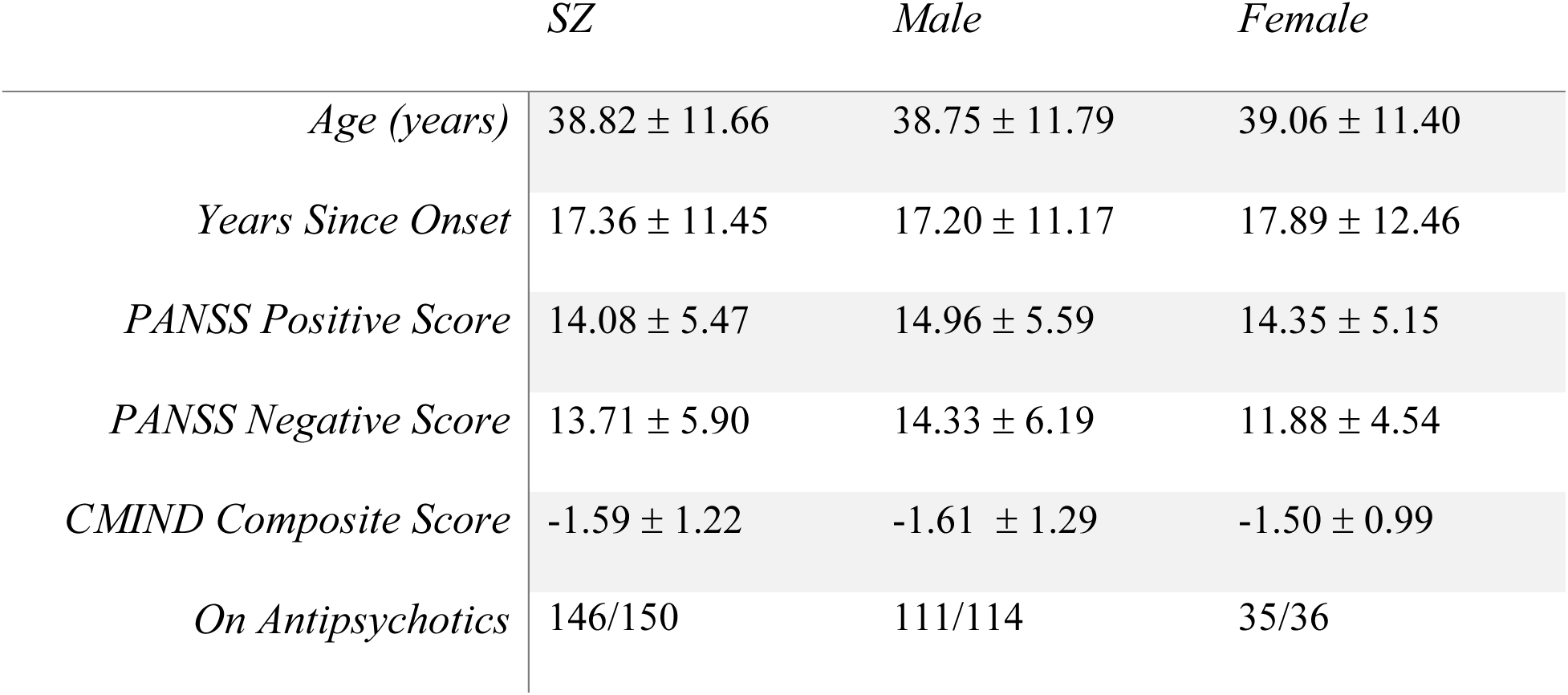
Demographic description of the SZ group.

|  | *SZ* | *Male* | *Female* |
| --- | --- | --- | --- |
| *Age (years)* | 38.82 ± 11.66 | 38.75 ± 11.79 | 39.06 ± 11.40 |
| *Years Since Onset* | 17.36 ± 11.45 | 17.20 ± 11.17 | 17.89 ± 12.46 |
| *PANSS Positive Score* | 14.08 ± 5.47 | 14.96 ± 5.59 | 14.35 ± 5.15 |
| *PANSS Negative Score* | 13.71 ± 5.90 | 14.33 ± 6.19 | 11.88 ± 4.54 |
| *CMIND Composite Score* | -1.59 ± 1.22 | -1.61 ± 1.29 | -1.50 ± 0.99 |
| *On Antipsychotics* | 146/150 | 111/114 | 35/36 |

Resting state fMRI (rsfMRI) data were collected with 3-Tesla MRI scanners with a repetition time (TR) of 2 seconds, voxel size of 3.44 x 3.44 x 4.00 mm, a slice gap of 1 mm, and a total of 162 volumes (∼ 5 minutes). Subjects were instructed to keep their eyes closed during the resting state scan but not to fall asleep. Preprocessing included brain extraction, slice-timing and motion correction steps. Preprocessed data were then registered into structural MNI space, resampled to 3 mm^3^ isotropic voxels, and spatially smoothed using a Gaussian kernel with 6 mm full-width at half-maximum (FWHM) on a per-subject basis. The first ten timepoints were trimmed from the time course and all voxel time courses were subsequently z-scored. Finally, we applied spatially constrained ICA (scICA) using the NeuroMark pipeline (Du et al., 2020) in the GIFT toolbox (http://trendscenter.org/software/gift & (Iraji et al., 2021)) to extract subject-level spatial maps for each of the 53 intrinsic connectivity networks (ICNs) of the NeuroMark_fMRI_1.0 template (http://trendscenter.org/data), as well as the respective activation time courses for each of the ICNs.

Structural MRI (sMRI) data were preprocessed using statistical parametric mapping (SPM 12) under the MATLAB 2019 environment. Structural images were segmented into grey matter, white matter, and cerebral spinal fluid (CSF) using a unified segmentation approach followed by modulation with the Jacobian of the transform (Penny et al., 2006), resulting in outputs as grey matter volume (GMV). Finally, the GMV maps were smoothed using a 3D Gaussian kernel with FWHM = 6 mm.

### 2.2 Filter-Banked Connectivity

As described in (Faghiri et al., 2021), the SWPC centered at each time point, *r_x,y_(t)*, for two time series *x(t)* and *y(t)* can be approximated by the following convolution, *g_x,y_(t)*:

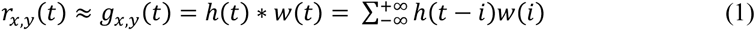

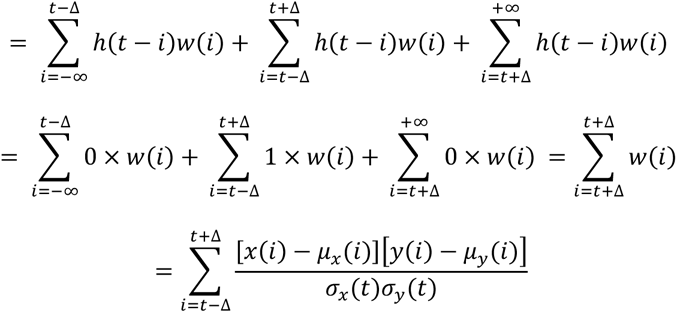

Where:

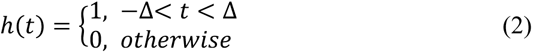

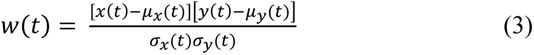

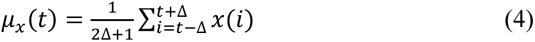

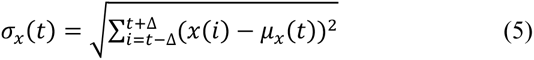

Per the system and signal theorem (Oppenheim & Schafer, 2010) the *g_x_*_,*y*_(*t*) series, and thus the SWPC that it approximates, can be seen as the output of a system with an impulse response *h*(*t*) (a rectangular window) and input an of *w*(*t*) (connectivity time series), resulting in a low-pass signal examining the low frequency range of *w(t)* (Fig. 1A). In the FBC approach, the *h(t)* of the SWPC formulation is replaced with a filter bank, i.e., an array of systems used to filter a time series into different frequency bands, usually non-overlapping spanning the entire frequency spectrum of the series. Each filter in the filter bank is defined by a response function *h_m_(t)*, where *m* is the filter index, resulting in *M* time series, each estimating the connectivity in a given frequency band (Fig. 1B). The filter bank design is fully flexible and can be tailored to best accommodate the spectral range of the data or aims of the analysis at hand. Thus, the FBC of two time series *x(t)* and *y(t)*, *r_m,x,y_(t)*, is defined as:

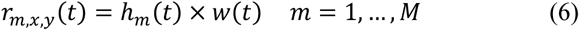

**Figure 1.**
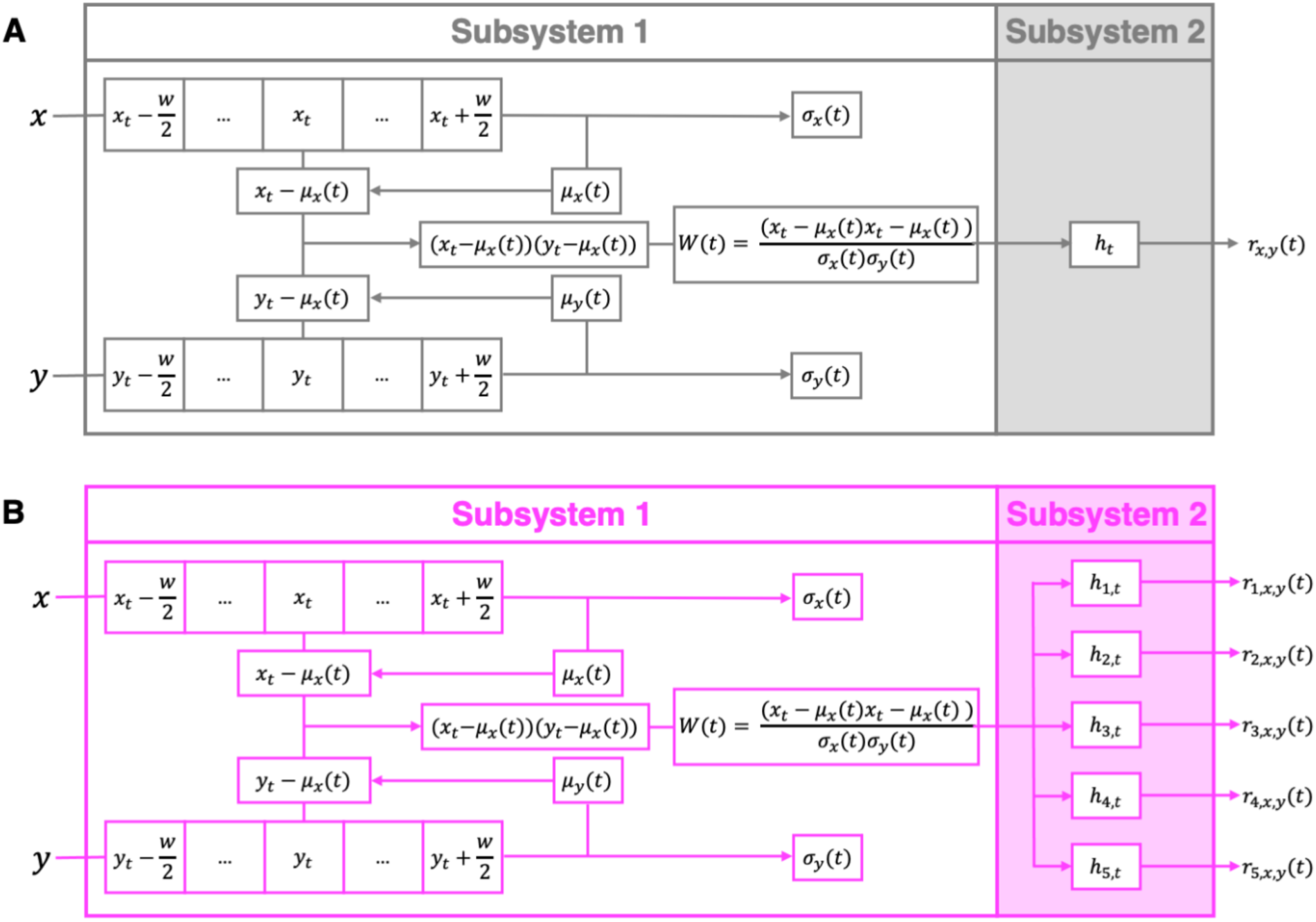
SWPC (A) and FBC (B) systems. While subsystem 1 is shared between both SWPC and FBC, in subsystem 2, SWPC uses a low-pass filter to examine the low-frequency range of w(t) (A) while FBC uses an array of filters to examine connectivity across various frequency bands (B). Thus, FBC is more flexible as it effectively combines both sFNC and dFNC, does not make assumptions about the connectivity frequency, and effectively spans a wide range of window sizes.

We calculated *w(t)* using a window *w* = 10 TR (22 s) for each pair of ICNs, resulting in 1378 (53 × (53 – 1)/2) *w(t)* time series. The filter bank was applied to each *w(t)* series separately using a forward-backward approach to achieve zero-phase filtering. We designed our filter bank to contain 10 Chebyshev type-2 infinite impulse response filters, the orders of which were obtained using `cheb2ord` as implemented in MATLAB to obtain at least 30 dB attenuation in the stopband and at most 3 dB in the passband (Rabiner & Gold, 1975). The 10 filters evenly cover the full frequency spectrum of the fMRI time series (0.00 – 0.25 Hz) as follows:

- Band 1: 0.000–0.025 Hz
- Band 2: 0.025–0.050 Hz
- Band 3: 0.050–0.075 Hz
- Band 4: 0.075–0.100 Hz
- Band 5: 0.100–0.125 Hz
- Band 6: 0.125–0.150 Hz
- Band 7: 0.150–0.175 Hz
- Band 8: 0.175–0.200 Hz
- Band 9: 0.200–0.225 Hz
- Band 10: 0.225–0.250 Hz

We applied k-means clustering to the FBC series stacked across all subjects and frequency bands to identify distinct states with unique connectivity signatures and spectral occupancy across frequency bands. Finally, we computed the subject-level mean connectivity for each state and concatenated them along with state-wise spectral occupancy to define the feature space for the fMRI modality for each subject. (Fig 2).

**Figure 2.**
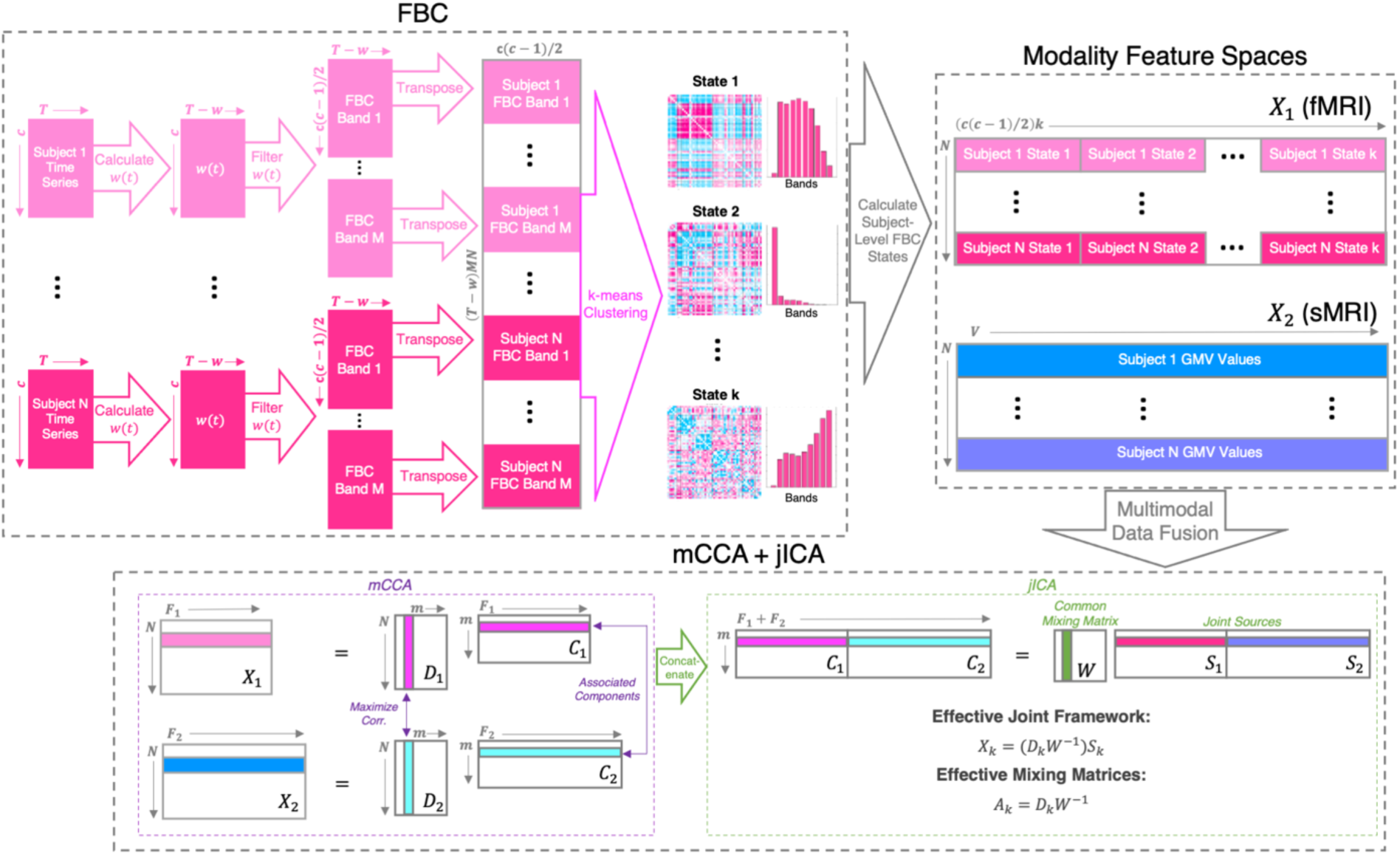
Filter-banked fusion pipeline. We applied FBC to fMRI data to extract subject specific FBC states, then applied the mCCA + jICA framework to extract joint sources, S1 & S2, from the fMRI FBC states (X_1_) and sMRI grey matter volume (X_2_).

### 2.3 Data Fusion: mCCA + jICA Framework

We used mCCA + jICA to perform fusion of the feature spaces generated from two imaging modalities, fMRI (processed using FBC) and sMRI (GMV maps) (Fig. 2). The mCCA + jICA framework is defined under the assumption that a multimodal dataset, *X_k_*, is a linear mixture of *m* sources (*S_k_*) mixed by non-singular matrices (*A_k_*), here, *k* = (1,2). The framework consists of two phases. The first mCCA phase begins with a dimensionality reduction step on the feature space of both modalities using principal components analysis (here PC = 100). Next, the canonical variates, *D_k_*, are estimated by maximizing the sum of squared correlations cost in *m* columns of the canonical variates (here *m* = 10). Last, the canonical correlation coefficients (CCCs) are estimated as association maps, *C_k_*, by inverting the *X_k_* = *D_k_C_k_* model.

In the second phase of the joint framework, the estimated CCCs are concatenated [*C_1_*, …, *C_k_*] and input into the jICA linear mixing model, *[C_1_, …, C_k_] = W[S_1_, …, S_k_]*. This decomposition reveals *m* maximally independent joint sources *S*, each of which contains a concatenation of co-varying modality-specific components. Thus, the effective mCCA + jICA framework can be defined as *X_k_ = (D_k_W^-1^)S_k_*, where the modality-specific mixing matrices are defined as *A_k_ = D_k_W^-1^*. Further details can be found in (Abrol et al., 2017; Sui et al., 2011, 2013).

## 3 Results

### 3.1 Filter-Banked Connectivity States

Using the elbow criterion on the within-cluster distance, we found six clusters to be optimal in the k-means analysis, each corresponding to a distinct connectivity state with a unique connectivity signature and spectral occupancy across the 10 frequency bands (Fig. 3). These states can be broadly split into low-pass (states 1-2), band-pass (states 3-5), and high-pass (state 6) frequency ranges. Significant group differences in subject-level fractional occupancy (i.e., percentage of all time points across all bands assigned to that state) were found in all six states. For example, we found the two low-frequency states could be further separated into a control-dominant (state 1) low-frequency state and a SZ-dominant (state 2) low-frequency state. The control-dominant low-frequency state was highly organized and characterized by integration of a sensory block comprised of the sensorimotor, visual, and auditory subdomains, which exhibited strong positive connectivity within the block and strong anticorrelations between the sensory block and the rest of the brain. In contrast, the SZ-dominant low-frequency state exhibited less complex functional organization, as it was characterized mainly by inter-domain connectivity only, as well as comparatively lower connectivity strength overall. At the other end of the spectrum, we found that the SZ group spent significantly more time in the high-frequency state 6 then the control group, which was consistent with the results reported in the original FBC work (Faghiri et al., 2021). This high-frequency state was marked by interesting cross-domain synchrony between the subcortical domain and the auditory and sensorimotor domains, as well as between the default mode domain and the cerebellum, with additional strong anticorrelation observed between these two blocks of cross-domain synchrony (i.e., SC/AUD/SM block anticorrelated with DM/CB block). Finally, we found that the two states with the lowest SZ fractional occupancy (states 1 and 3) have nearly opposing connectivity signatures, both marked by strong correlation (or anti-correlation) within the sensory domain block as well as strong anticorrelation (or correlation) between the sensory domain block and all other functional domains, with the strongest FC antagonism seen between the sensory block and the subcortical domain.

**Figure 3.**
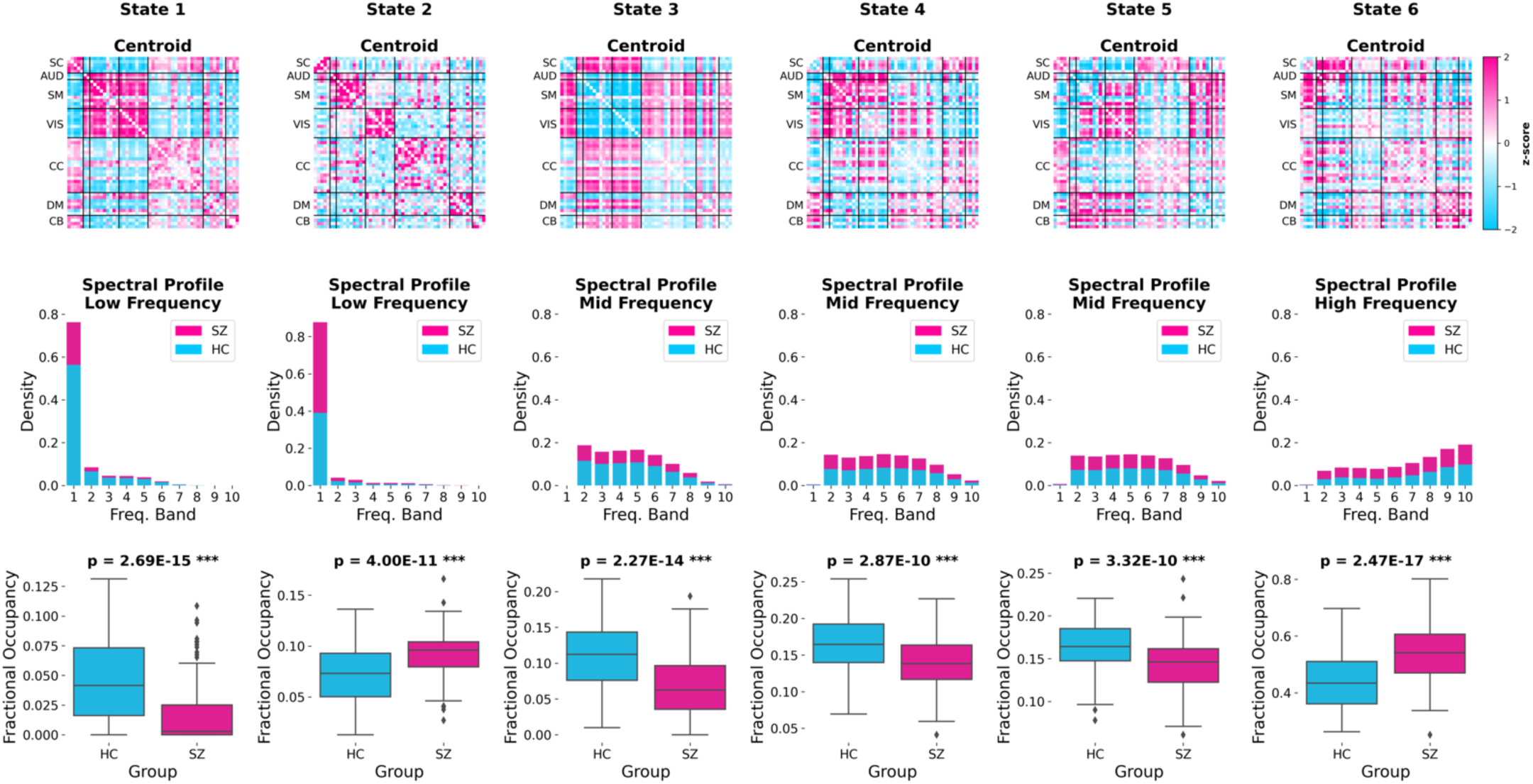
Summary of FBC States. State centroids shown as z-scored connectomes in the top row, spectral profiles are shown as stacked fractional occupancy histograms across the ten frequency bands in the middle row, and group-level state occupancy is shown in the boxplots on the bottom row. States 1-2 are predominantly identified in low-frequency bands, states 3-5 are predominantly identified in mid-frequency bands, and state 6 is predominantly identified in high-frequency bands. All p-values corrected for multiple comparisons (FDR).

### 3.2 Joint Sources

Of the ten joint sources (determined by the chosen model order) that were extracted, two had significant group differences (after FDR correction) in loadings for both the structural and functional components of the joint source. Summaries of these joint sources are presented in the following sections.

#### 3.2.1 Joint Source 1

A summary of the first joint source is shown in Fig. 4. The structural component for this joint source showed peaks in grey matter volume alterations in the middle temporal gyrus, precentral gyrus, insula, right inferior frontal gyrus, left inferior parietal lobule and anterior cingulate cortex (Fig. 4B). The linked functional component of the joint source showed frequency-specific connectivity patterns across each of the FBC states, however significant edges involving the subcortical domain were commonly identified across all six states. All significant edges (|z| > 2.5) across all states are shown in Fig. 4A, but here we highlight a few patterns of interest. In the low-frequency states, the functional components contained opposing patterns of connectivity within the subcortical domain, as well as between the subcortical and sensorimotor domains, where the control-dominant state 1 component contained anticorrelation within the subcortical domain and positive correlation between the subcortical and sensorimotor domains while the SZ-dominant state 2 component was marked by within-domain subcortical synchrony and cross-domain anticorrelation between the subcortical and sensorimotor networks.

**Figure 4.**
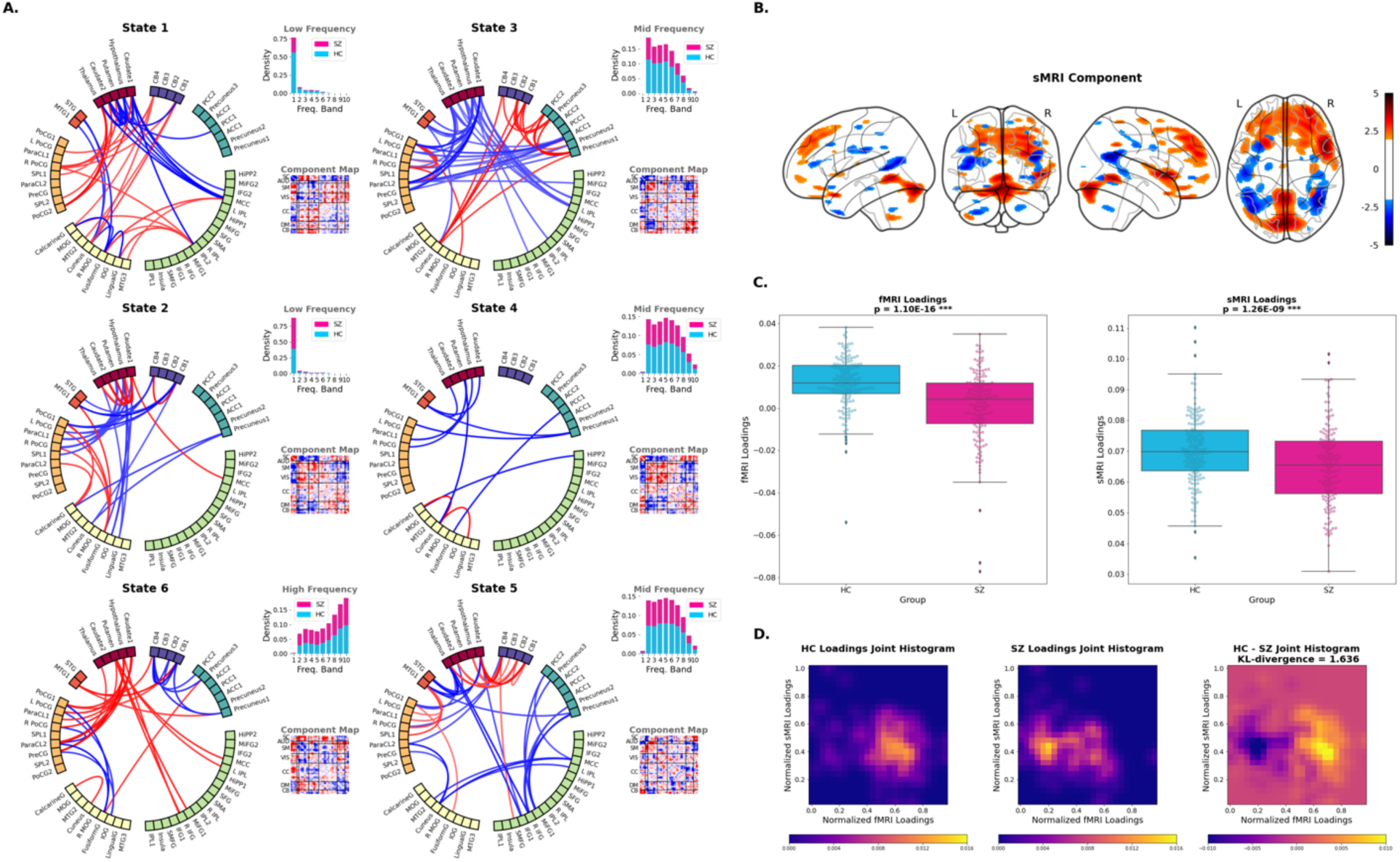
Summary of Joint Source 1. (A) Significant edges (i.e., functional connections with connectivity strength |*z*| ≥ 2.5) in each FBC state for the functional component of the joint source. Colors of nodes show network affiliation and colors of edges denote positive (red) or negative (blue) connectivity. Stacked bar graphs of the spectral profiles as well as the full component maps as connectome matrices are also shown for each state. (B) Spatial map of the significant (|*z*| ≥ 2.5) regions of the structural component of the joint source. (C) Loading parameters show strong group differences for both the functional (p = 1.10×10^-16^) and structural (p = 1.26×10^-9^) components. (D) Joint histograms of the fMRI and sMRI loadings show that the relationships between the structural and functional components of the joint source are strongly group-specific (Kullback-Leibler divergence = 1.636).

Interestingly, the components of the two lower-frequency control-dominant states (1 and 3) also shared distinctive connectivity features–functional correlation between cerebellar regions and the cuneus in the visual domain as well as anticorrelation between subcortical regions and regions in the cognitive control domain, namely the middle cingulate cortex and the left inferior parietal lobule. The SZ-dominant high-frequency state 6 component map showed an opposing pattern of strong positive correlation between the subcortical domain, specifically the precuneus, and the middle cingulate cortex and the left inferior parietal lobule within the cognitive control domain. In addition, the state 6 component map was marked by strong positive correlations between the subcortical domain and the sensorimotor domain, which mirror patterns from the state 1 component, within-domain anticorrelation of the subcortical domain, which mirror patterns seen in state 2, as well as strong anticorrelations between the cerebellum and default mode domains, which are not seen in any other state component of the joint source.

We found significant group differences in the loading parameters (derived from mixing matrix *A_k_*) for both the functional (*p* = 1.10×10^-16^) and structural (*p* = 1.26×10^-9^) components (Fig. 4C), with the SZ group exhibiting significantly lower loadings than the control group in both cases, indicating the SZ group had significantly reduced expression of the structural and functional patterns represented by the respective structural and functional component maps. There was also a significant correlation (*r* = 0.416; *p* = 1.01×10^-13^) between the loading parameters of the structural and functional components; however, the joint histograms of the structural and functional loadings in Fig. 4D suggest the relationship between the structural and functional components is more complex than a simple linear correlation, and in fact, this relationship differs significantly between the SZ and control groups, as evidenced by the Kullback-Leibler divergence (KLD) = 1.636 between the two group joint histograms.

#### 3.2.1 Joint Source 2

A summary of the second joint source is shown in Fig. 5. The structural component map for this joint source contained a pattern of higher GMV in regions within the motor cortex, as well as lower GMV within the cerebellum (Fig. 5B). Again, the linked functional component of the joint source contained unique connectivity patterns within each of the frequency-specific states, however functional connections involving the sensorimotor and cerebellar domains were prominent across all FBC state functional component maps. All significant edges (|z| > 2.5) across all states are shown in Fig. 5A, but here we highlight a few patterns of interest. The low-frequency state 1 functional component was highly organized and mostly defined by strong functional integration (i.e., positive connectivity) between the cerebellar domain with nearly all regions of the sensorimotor and visual domains, as well as anticorrelation of sensorimotor networks with regions in the cognitive control domain, specifically the supplementary motor area, inferior frontal gyrus and the superior medial frontal gyrus. Conversely, the SZ-dominant low-frequency state 2 showed largely opposing patterns of cerebellar connectivity, characterized mainly by anticorrelation between the cerebellum and both sensorimotor and visual regions. State 2 also showed strong within-domain connectivity in the visual domain, as well as some positive correlation of the visual domain with the superior parietal lobule and postcentral gyrus in the sensorimotor domain. The mid-frequency state 3 was dominated by connections involving regions within the sensorimotor domain to nearly all other domains in the brain, with the notable exception being the absence of connections between the sensorimotor and cerebellar domain above our significance threshold. The mid-frequency state 4 functional component included a connectivity pattern that was not seen in any of the other state components–strong positive correlations between the visual domain and several regions in the cognitive control domain, mainly encompassing the inferior parietal lobule, the middle frontal gyrus and inferior frontal gyrus, as well as some negative correlations between visual domain networks and the hippocampus, also of the cognitive control domain. Lastly, the SZ-dominant high-frequency state 6 was defined by strong anticorrelations of the subcortical networks with sensory domains including auditory, sensorimotor and visual domains, paired with strong integration within the sensorimotor domain and between the sensorimotor and auditory domain. There was no significant integration of the sensorimotor and visual domains in the state 6 functional component, however both the sensorimotor and visual domains did exhibit strong positive correlation with specific cognitive control networks, the former with the supplemental motor area and the latter with the superior frontal gyrus.

**Figure 5.**
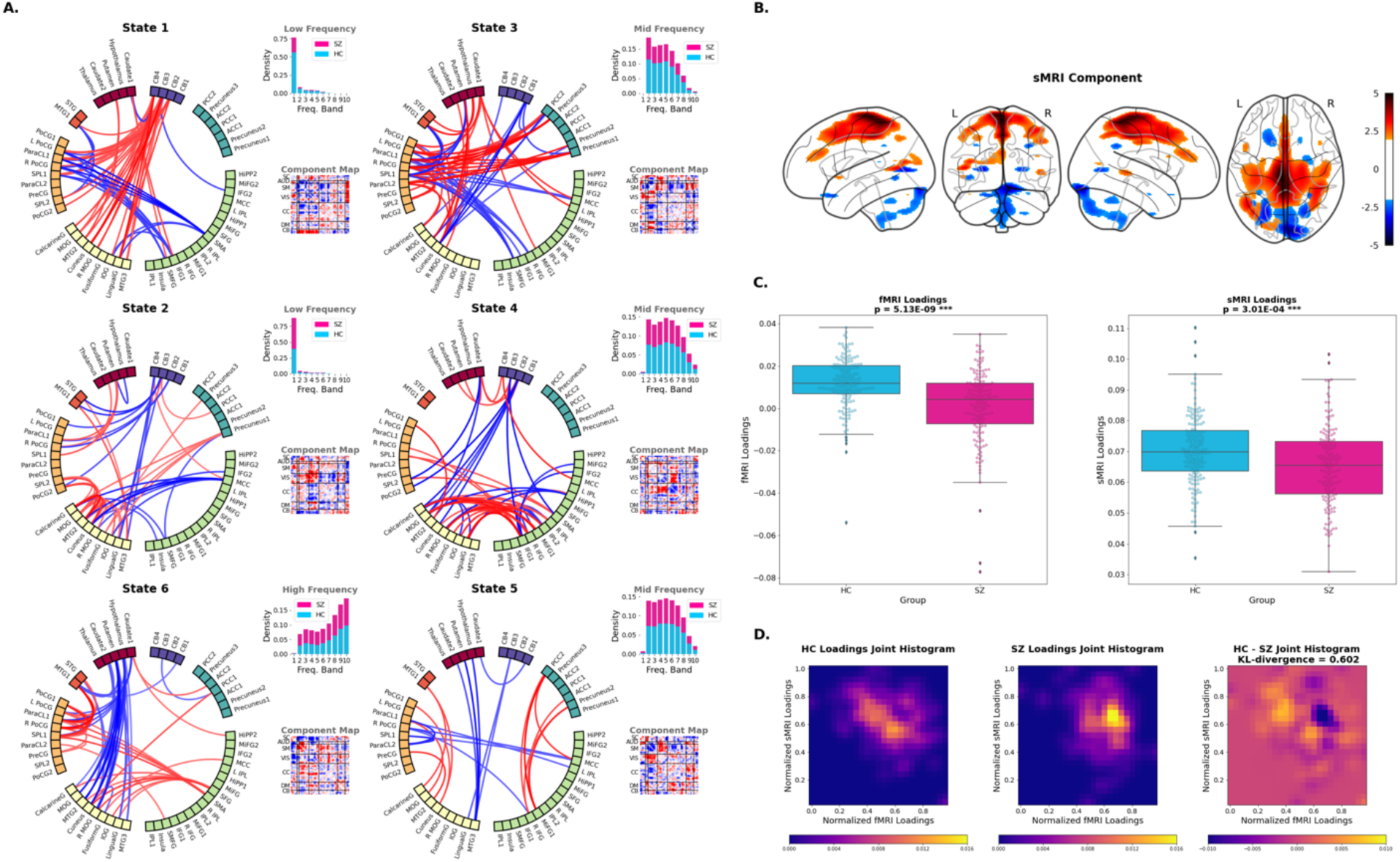
Summary of Joint Source 2. (A) Significant edges (i.e., functional connections with connectivity strength |*z*| ≥ 2.5) in each FBC state for the functional component of the joint source. Colors of nodes show network affiliation and colors of edges denote positive (red) or negative (blue) connectivity. (B) Spatial map of the significant (|*z*| ≥ 2.5) regions of the structural component of the joint source. (C) Loading parameters show strong group differences for both the functional (p = 5.13×10^-9^) and structural (p = 3.01×10^-^) components. (D) Joint histograms of the fMRI and sMRI loadings show that the relationships between the structural and functional components of the joint source are strongly group-specific (Kullback-Leibler divergence = 0.602).

Similar to the first joint source, we found significant group differences in the loading parameters for both the functional (*p* = 5.01×10^-9^) and structural (*p* = 2.99×10^-4^) components (Fig. 5C), with the SZ group again exhibiting significantly lower loadings than the control group in both cases, indicating reduced overall expression of these functional and structural patterns within the SZ group. We again found a significant correlation (*r* = 0.474; *p* = 9.15×10^-19^) between the loading parameters of the structural and functional components; however, the joint histograms of the structural and functional loadings in Fig. 5D provide evidence that again the relationship between the structural and functional components is more complex than a simple linear correlation. We find a high density of SZ subjects fall within a small region of the joint histogram, and a more diffuse dispersion of control individuals in their group joint histogram that suggests an anticorrelation relationship between structural and functional loadings within the controls. We found the KLD = 0.602 between the two group joint histograms.

## 4 Discussion

In this work, we investigated the relationship between frequency-specific patterns of functional connectivity and structural measures of GMV to elucidate key structure/function relationships implicated in schizophrenia. Specifically, we utilized the newly proposed FBC approach to estimate dFNC across ten non-overlapping frequency bands and ultimately derive six distinct FBC states, each defined by its own unique frequency range. We then utilized the mCCA + jICA symmetric multimodal data fusion framework to identify hidden linkages between the connectivity patterns of these frequency-specific FBC states and grey matter volume maps from sMRI in the form of jointly co-varying functional and structural components, here called joint sources.

The FBC analysis identified six connectivity states characterized by unique spectral profiles as well as connectivity patterns. The most interesting group differences in fractional occupancy of these states were found at the lowest and highest frequency ranges. Of the two low-frequency FBC states, one was defined by strong synchrony within the somatosensory block (sensorimotor, auditory, and visual domains) that was anticorrelated with the rest of the brain (most strongly with subcortical regions), and was primarily occupied by healthy controls, and the other was characterized by strictly within-domain synchrony as well as overall lowered connectivity strength, which was significantly dominated by SZ occupancy. This result is in line with previous works that report generalized lower connectivity in SZ compared with controls (Bluhm et al., 2007; Dong et al., 2018; Erdeniz et al., 2017; Liang et al., 2006; Lynall et al., 2010; Meda et al., 2012; Skudlarski et al., 2010) and conforms with the dysconnectivity hypothesis of SZ (Friston & Frith, 1995), which posits that dysfunctional integration of brain networks and generally disconnected or misconnected neural circuitry might contribute to the pathophysiology of SZ. The identification of this dichotomy in connectivity strength and functional organization between SZ subjects and controls in the low frequency range in our results is not unexpected, as this phenomenon has been reported in studies of largely static FNC or SWPC-based dynamic FNC, which we have established miss the higher frequency states the FBC approach is capable of extracting (Faghiri et al., 2021).

We also identified one state in the high frequency spectral range, which had the highest SZ occupancy of all six states, as well as the most significant group difference in occupancy between SZ and HC. This result is in line with the prior FBC work which found that individuals with SZ spend more time in high frequency states than control individuals (Calhoun et al., 2008; Faghiri et al., 2021; Turner et al., 2013). (Yaesoubi et al., 2017) similarly reported SZ subjects were more likely to occupy the highest frequency state; however their method was based on frequency analysis in the *activity* domain rather than the *connectivity* domain like in the FBC approach, which resulted in vastly different connectivity profiles for the high frequency states between their work and ours. This discrepancy again underscores the fact that the relationship between the activity and connectivity domains is not clear. There is evidence from fMRI studies of increased power spectra of certain ICNs (e.g., default mode) at higher frequencies in individuals with SZ (Calhoun et al., 2011; Garrity et al., 2007) as well as EEG/MEG studies that show an association between aberrant neural oscillations in the high frequency beta and gamma bands and SZ (Moran & Hong, 2011; Roach et al., 2013; Tan et al., 2013; Uhlhaas & Singer, 2013). While these studies also apply frequency-based analyses on the activity domain of the functional neuroimaging signal, this convergence of evidence across a range of methodologies heavily implicates altered high frequency brain function in SZ.

The role of subcortical (particularly thalamic) and somatosensory connectivity in SZ has often been reported in the literature (Anticevic et al., 2014; Cao et al., 2022; DeRamus et al., 2022; Ferri et al., 2018; Skåtun et al., 2017, 2018; Welsh et al., 2010). Sensory regions including auditory, visual, and sensorimotor networks have been implicated in possible “bottom-up” processes that may contributing to a range of emotional and cognitive symptoms associated with SZ (Javitt, 2009; Revheim et al., 2014). Furthermore, the sensory gating hypothesis (Cromwell et al., 2008) suggests the process the brain uses to filter and assign importance to external stimuli is abnormal in SZ, strongly implicating both thalamic dysfunction, as well as aberrant functional synchronization between the thalamus and frontal/somatosensory networks. A recent pharmaco-FMRI study using the NMDA receptor (NMDAR) antagonist, ketamine, implicated NMDAR hypofunction as a mediator of this thalamo-cortical dysconnectivity pattern across the illness course of schizophrenia, including the psychosis-risk syndrome that sometimes progresses to full schizophrenia (Abram et al., 2022). Though there is mounting evidence that somatosensory/subcortical dysfunction plays a role in SZ pathophysiology, conflicting results have been published on the nature of this dysfunction–some reporting higher connectivity (or hyperconnectivity) between subcortical and sensory regions (Damaraju et al., 2014; Fu et al., 2018; Yaesoubi et al., 2017; D. Zhang et al., 2012), while others report lower connectivity (or hyperconnectivity) between these networks (Skåtun et al., 2017; Welsh et al., 2010; Y. Zhang et al., 2021). In our work, three of our six FBC states are characterized by strong connectivity (defined by both strongly positive or strongly negative correlations) between subcortical and somatosensory regions: states 1, 3, and 6. Interestingly, the states at the lower end of the frequency spectrum (states 1 and 3) with this functional relationship are the states in which we observed higher fractional occupancy of control individuals paired with the lowest fractional occupancy of SZ individuals among all the states, while the high frequency state 6 that shows evidence for strong subcortical-sensory synchrony was marked by significantly higher SZ occupancy. Thus, our results suggest that in SZ subcortical-sensory connectivity may be weaker or absent at lower frequencies while strong synchrony between these regions may exist when higher frequency functional connectivity fluctuations are considered.

We identified two joint sources that exhibited significant group differences in both structural and functional component loadings, indicating these joint sources do indeed encode structure-function relationships that are frequency-dependent and relevant to SZ. The first joint source implicated regions in the middle temporal gyrus, precentral gyrus, insula, right inferior frontal gyrus, left inferior parietal lobule and anterior cingulate cortex. This component closely resembles the combinations of two structural components found to have the highest effect size between SZ and control groups via source based morphometry (SBM) analysis of structural MRI data alone (Gupta et al., 2015, 2017). Inspection of group differences in loading parameters revealed SZ subjects had significantly lower loading values than the controls, indicating a generally weaker expression of the component pattern of GMV in these areas related to SZ. The related functional component shows functional connectivity patterns that are clearly frequency-specific across the six states, and we observed that many of the significant edges across the state-level functional components involve subcortical-somatosensory connections. Opposing subcortical-sensory connectivity patterns were identified in the two low frequency states, with the SZ-dominant state 2 defined by synchrony within the subcortical domain but anticorrelation between subcortical/sensorimotor, while the control dominant state 1 was defined by anticorrelation within the subcortical domain as well as subcortical-sensorimotor synchrony.

Importantly, the subcortical-sensorimotor synchrony was also a hallmark of the high-frequency and SZ-dominant state 6 component, further indicating that there may be frequency-based modulation of subcortical-somatosensory connectivity contributing to the functional pathophysiology of SZ.

In the second joint source, we identified structure/function linkages between GMV in the motor cortex and cerebellum with frequency-specific functional connections within the same domains. Lower GMV in the cerebellum and its link to the cerebellar motor module (connection between the cerebellum to the cortical sensorimotor network) has been previously reported in SZ (He et al., 2018). Again, the functional components of the low frequency states show opposing connectivity signatures, where the low-frequency state 1 functional component was highly organized and mostly defined by strong functional integration between the cerebellar domain with nearly all regions of the sensorimotor and visual domains while the SZ-dominant low-frequency state 2 was characterized mainly by anticorrelation between the cerebellum and both sensorimotor and visual regions. Evidence for stronger cerebellar-somatomotor connectivity in SZ compared to HC has been reported (Shinn et al., 2015), and our results suggest this hyperconnectivity linked to motor/cerebellar GMV alterations exists mainly at low-to-mid frequency ranges. In fact, the high frequency functional component (state 6) contains no cerebellar-sensorimotor linkages, but rather is largely characterized by subcortical-sensory edges, further suggesting the importance of these functional connections at high frequencies.

Beyond the structural and functional components themselves, our results provide evidence that the relationship between the identified structural and functional patterns differs between individuals with SZ and controls. Significant positive correlations were found between the structural and functional loading parameters of both joint sources (r = 0.416, p = 2.02×10^-19^; *r* = 0.474; *p* = 9.15×10^-19^, respectively). However, additional analysis of the joint sMRI/fMRI loading parameters revealed that the relationships between the structural and functional components required a more nuanced interpretation across our diagnostic groups than just linear correlation. For both joint sources there existed a significant difference in density and distribution of subjects within the joint histogram between the SZ and control groups, indicating that the association between the structural and functional components varied in a manner that was not completely linear. This was especially evident for the first joint source, where the KLD between groups was larger than that of the second joint source (KLD = 1.64 vs 0.60, respectively), indicating the distributions of structural/functional loadings between patients and controls were even further apart. More work is needed to disentangle these exact relationships further.

Many of the regions identified in our joint sources have been previously implicated in SZ, supporting the results of prior work across both unimodal and multimodal methodologies. However, our investigation is distinguished from these prior studies as it is, to our knowledge, the first multimodal study to include frequency information, specifically frequency in the *connectivity* domain rather than the *activity* domain, in the fMRI feature space. Thus, our results help shed new light on the underlying nature of structure/function relationships characteristic of the SZ brain. For instance, our results suggest that cortico-subcortical connections, specifically those between subcortical and somatosensory regions, are of particular importance in high-frequency ranges and do indeed co-vary with structural alterations in GMV across a variety of brain regions in SZ. These and other linkages reported here may have been missed, or the nature of the functional oscillations in connectivity not fully understood, as the typical SWPC method for estimating dFNC has been shown to miss high-frequency states like state 6 in our results (Faghiri et al., 2021).

Our study has some limitations that should be considered. First, our analysis was performed on a single dataset with a sample size of N = 310, which can be considered large compared to classic imaging studies where only tens of subjects were scanned but can also be seen as relatively small compared to publicly available imaging datasets where sample size can reach 1000+ subjects. Replication of these results in an independent dataset should be a focus of future work. Second, the fMRI data used to estimate our FBC states has a relatively low temporal resolution of TR = 2 sec. Since the available frequency range is tied directly to the temporal resolution (i.e., sampling rate) of the data, it would be beneficial to repeat our analysis in data with higher temporal resolution (e.g., TR < 1 sec.) to expand the frequency range within which the FBC states can be estimated. Considering the strong evidence of the importance of the very high frequency connectivity states as a functional component of SZ, we believe it will be extremely beneficial to explore these high frequency ranges more granularly as higher temporal resolution image acquisitions become more readily available. The relatively short acquisition time of our data (∼5 minutes) could also be considered as a potential limitation, and future work in this space may focus on replicability of our findings in longer or repeated scans. As mentioned frequently throughout our report, the key novelty of the FBC approach is its ability to apply time-frequency analysis directly in the connectivity domain rather than the activity domain. Recent work has focused on the nature of the linkage between activity and connectivity domains, and even provides evidence that this relationship may vary for HC and individuals with SZ (Fu et al., 2018, 2021). Future work may focus on a combined data fusion approach in the context of linking activity and connectivity together with structure. Future studies may also choose to treat each frequency-specific FBC state as a separate modality within the fusion architecture, rather than concatenating all the states into a single fMRI modality vector per subject. Such a study design would allow for more flexible linkages between each state and the structural components and add an opportunity for an additional layer of investigation and interpretation. A series of studies (Clementz et al., 2022) have shown that there is significant overlap between the structural and functional brain abnormalities reported in schizophrenia and those seen in psychotic bipolar and schizo-affective disorders. Thus, claims of specificity to schizophrenia of the findings reported here remain to be demonstrated. Finally, the interpretation of our results should be considered in the context of the history of antipsychotic and other medication in the SZ group.

In conclusion, our results suggest there is a frequency-specific functional component of the structure/function relationship underlying the pathophysiology of SZ, particularly at the lowest and highest connectivity frequencies.

## 5 Acknowledgements

We would like to acknowledge funding from the National Institute of Mental Health (Grant/Award Number: R01MH123610), the National Science Foundation (Grant/Award Number: 2112455), and author JMF’s VA Senior Research Career Scientist Award (Grant/Award Number: 1IK6CX002519).

## 6 Data/Code Availability

Details on the availability of the FBIRN dataset used as our discovery dataset can be found at https://www.nitrc.org/projects/fbirn/. The code and network templates used for spatially constrained ICA, as well as the data fusion toolbox, are available at http://trendscenter.org/software.

## Notes

### Competing Interest Statement

The authors have declared no competing interest.

